# Time-Resolved Chemical Phenotyping of Whole Plant Roots with Printed Electrochemical Sensors and Machine Learning

**DOI:** 10.1101/2023.03.09.531921

**Authors:** Philip Coatsworth, Yasin Cotur, Atharv Naik, Tarek Asfour, Alex Silva-Pinto Collins, Selin Olenik, Laura Gonzalez-Macia, Tolga Bozkurt, Dai-Yin Chao, Firat Güder

## Abstract

Plants are non-equilibrium systems consisting of time-dependent biological processes. Phenotyping of chemical responses, however, is typically performed using plant tissues, which behave differently to whole plants, in one-off measurements. Single point measurements cannot capture the information rich time-resolved changes in chemical signals in plants associated with nutrient uptake, immunity or growth. In this work, we report a high-throughput, modular, real-time chemical phenotyping platform for continuous monitoring of chemical signals in the often-neglected root environment of whole plants: TETRIS (**<u>T</u>**ime-resolved **<u>E</u>**lectrochemical **<u>T</u>**echnology for plant **<u>R</u>**oot ***<u>I</u>****n-situ* chemical **<u>S</u>**ensing). TETRIS consists of screen-printed electrochemical sensors for monitoring concentrations of salt, pH and H_2_O_2_ in the root environment of whole plants. TETRIS can detect time-sensitive chemical signals and be operated in parallel through multiplexing to elucidate the overall chemical behavior of living plants. Using TETRIS, we determined the rates of uptake of a range of ions (including nutrients and heavy metals) in *Brassica oleracea acephala.* We also modulated ion uptake using the ion channel blocker LaCl_3_, which we could monitor using TETRIS. We developed a machine learning model to predict the rates of uptake of salts, both harmful and beneficial, demonstrating that TETRIS can be used for rapid mapping of ion uptake for new plant varieties. TETRIS has the potential to overcome the urgent “bottleneck” in high-throughput screening in producing high yielding plant varieties with improved resistance against stress.

## 1. Introduction

The development of stress resistant varieties of plants that can tolerate biotic (*e.g.,* fungi, bacteria, viruses) and abiotic (*e.g.,* salinity, drought, heat, cold, extreme pH) stresses, while still producing high yields, is crucial to maintaining and improving crop production. Conventional (including glasshouse- and field-based) phenotyping (assessment of, for example, physical form and visual appearance) to select for stress tolerant species and varieties typically only results in a single yield measurement for a set condition or growth season.^1^ Plants under stress, however, produce chemical signals that often vary in concentration or composition over time.^2^ For the accelerated development of stress resistant species, measuring the behavior of the plant in real-time under stressful, but realistic, simulated environmental conditions is, therefore, of great interest both in applied and fundamental research.

Simulated growth conditions (*e.g.,* laboratory or glasshouse) can be combined with analytical methods to quantify biometrics, improving the efficiency and accuracy of rapid and automated phenotyping. Among various analytical techniques, optical approaches have probably been used the most widely in the research community for continuous-time phenotyping of whole, living plants.

Depending on the phenotype in question, different optical methods can be employed, some of which produce signals with spatiotemporal fidelity. For example, infrared (IR) thermography, a method for non-invasively measuring IR heat signatures, has been used to monitor stomatal conductance (degree of stomatal opening used for estimating CO_2_ and H_2_O exchange^3, 4^) under conditions of salt stress, allowing estimation of robustness against salt stress.^5^ Positron emission tomography (PET), which spatially maps the radiation emitted by a compound, has been used to estimate resistance to salt stress by measuring the uptake and transport of ^22^Na radioisotope in salt-resistant and salt-sensitive plant species.^6, 7^ The capabilities of optical phenotyping can be expanded through the use of genetically encoded and engineered materials that can be introduced into plants. These materials can be used to monitor chemical signals such as pH or Ca^2+^ *via* detection of fluorescence to study stress responses from stressors such as insect feeding.^8–10^ Despite being a powerful approach for continuous-time phenotyping of whole plants in terms of analytical performance, optical sensing methods often require expensive and specialized equipment that may not be accessible or available to all laboratories.

Because of the cost and complexity of optical approaches, they are often difficult to scale for mass phenotyping at an industrial scale. Interference from plant tissues and external light sources may also affect monitoring.^11^

Electrochemical sensors for continuous-time phenotyping of whole plants are emerging as an alternative to optical methods.^12–16^ Insertable electrochemical sensors (sensors that are inserted into plant tissues, such as stems or leaves) are typically formed either of conductive electrodes or microfabricated field-effect transistors and have been used to measure the internal chemical environment of plant tissues. Insertable sensors can detect bursts of H_2_O_2_ resulting from pathogenic and UV light stresses in leaf tissue, enabling characterization of responses to these stressors.^17, 18^ Insertable sensors are, however, invasive (though minimally) and can potentially generate unintentional stress responses, reducing the signal-to-noise ratio for the signal of interest.^17–19^ Non-invasive electrochemical sensors for continuous-time phenotyping of whole plants come in different forms, including devices placed externally on plants for measuring gaseous analytes.^20–23^

Despite the critical importance of the roots as the main entrance of water and nutrients for the plant, application of non-invasive electrochemical sensors to the roots for real-time, whole plant phenotyping has been underexplored. Because of the importance of pH in the root microenvironment, Felle used a glass microelectrode to continuously measure the apoplastic pH of the root cortex of *Zea mays* (corn) to investigate the impact of various ions and compounds on pH regulation in the roots.^24^ Using the same system, Felle *et al.* studied pH signaling on both the root surface and in the leaf apoplast of *Hordeum vulgare* (barley) upon pathogenic inoculation of the roots, demonstrating time-dependent response of pH under pathogenic stress.^25^ Glass pH microelectrodes, although effective, are fragile instruments that could not be easily handled by non-expert users. Pt black/carbon nanotube-based microelectrodes were developed by McLamore *et al.* for the detection of phytohormone indole- 3-acetic acid (IAA) in *Zea mays* roots.^26^ IAA, which regulates response to abiotic stress, is taken into roots by both diffusion and transporters, and the microsensor demonstrated influx occurred at different amounts at different locations around the root. Influx also showed an oscillatory pattern over the scale of minutes, with different mutants producing responses with different amplitudes and time periods. Although this work illustrates the precision microelectrodes afford, they are less useful for revealing whole-plant behavior and share the same fragility limitations as the glass electrodes by Felle *et al.*.^24, 25, 27^ These systems are also not compatible with mass fabrication, and hence continuous-time phenotyping has not been adopted by the wider community to parallelize and automate phenotyping.

In this work, we report our whole-plant monitoring platform: TETRIS (**<u>T</u>**ime-resolved **<u>E</u>**lectrochemical **<u>T</u>**echnology for plant **<u>R</u>**oot ***<u>I</u>****n-situ* chemical **<u>S</u>**ensing) (**Figure 1**). TETRIS consists of three screen-printed low-cost sensing modules for the multiplexed measurement of concentrations of ions (by electrochemical impedance spectroscopy (EIS)), H_2_O_2_ and pH. Changes in pH and H_2_O_2_ levels can coincide with wounding and biotic stresses, such as pathogen infection.^28, 29^ Ions vary in size, charge and relevance to plants, where some are key nutrients and others are toxic heavy metals, and so tracking their uptake can be indicative of salt sensitivity or tolerance.^30, 31^ As the rate of uptake of ions depends on charge gradients, the simultaneous measurement of multiple analytes can elucidate the interactions between these different chemical species.^32, 33^ We have extensively characterized TETRIS and investigated the characteristics of uptake of ions in whole *Brassica oleracea acephala* (dwarf green curled kale) plants, a common crop plant, continuously and in real-time. We studied the significance of the substituent ions and their properties on the rate of uptake and investigated the effect of ion-channel blockers on the uptake of ions. Finally, using the library of data collected using TETRIS, we developed a machine learning model that can predict the rates of uptake of salts.

**Figure 1.**
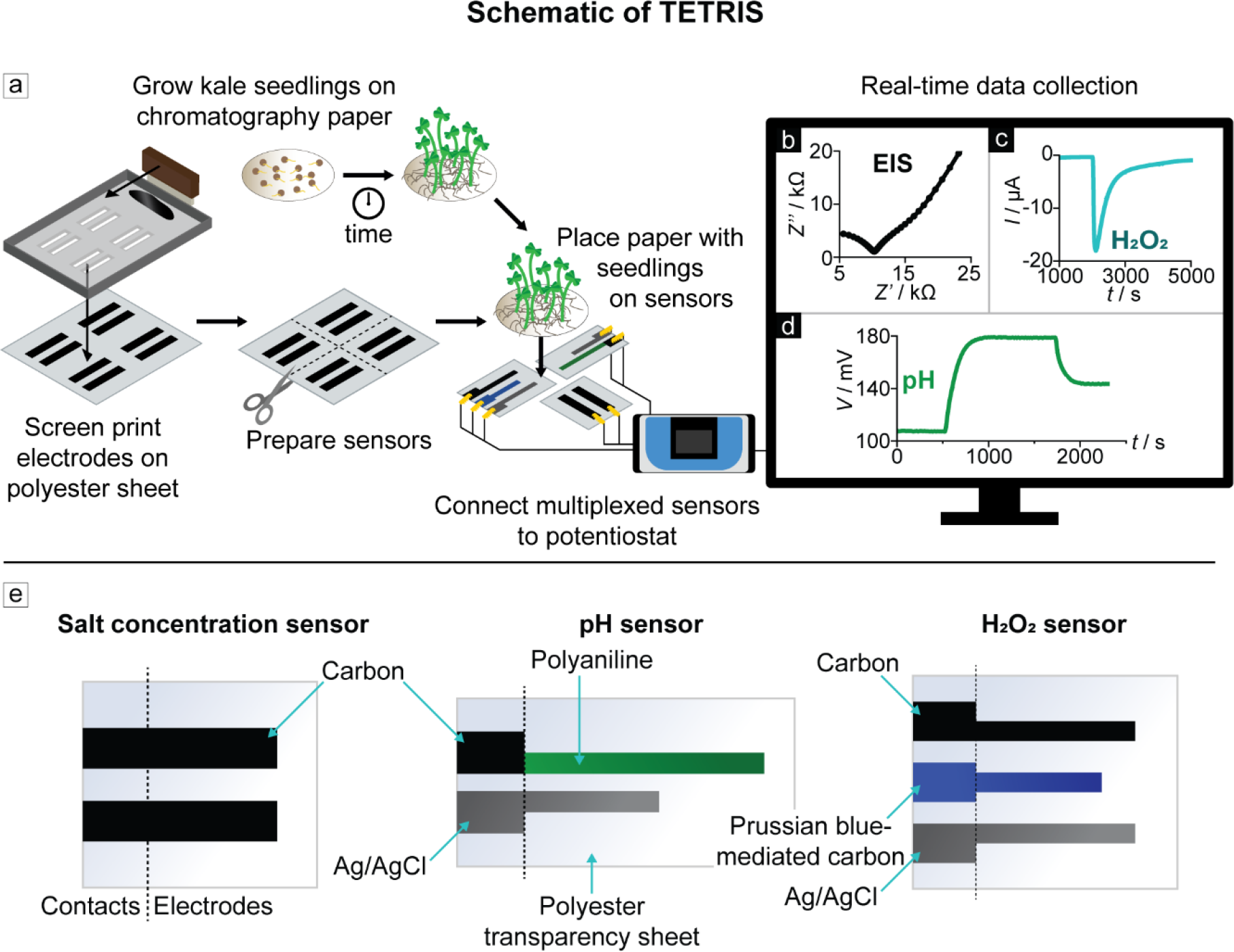
a) Schematic of TETRIS, showing sensor fabrication, growth of seedlings and recording of measurements using a standard laboratory potentiostat; **b)** Electrochemical impedance spectroscopy Nyquist plot of a paper disc with 30 kale seedlings (9-days-old); **c)** H2O2 sensing during addition of H2O2 (1 mM, 20 μL) to paper disc with KCl (1 M, 180 μL); **d)** pH sensing during addition of H2SO4 and NaOH to deionized water; **e)** Design and materials of our salt concentration, pH and H2O2 sensors, illustrations not to scale.

## 2. Results and discussion

### General experimental setup of TETRIS

To measure the chemical environment of the roots of plants, we developed a sensing platform, TETRIS, comprising a measurement chamber and disposable sensing module (Video S1). Sensors were created by screen-printing conductive ink onto plastic transparency sheets. Screen-printing is a technique that is low-cost, utilized industrially and does not require highly specialized equipment, meaning that transfer of these sensors from a laboratory to a production scale is reasonably straight-forward. The sensors are then individually cut (and for the pH sensor, electrochemically treated) and attached to a raised platform.

The module itself can hold a single sensor or multiple sensors in either stacked or lateral positioning to form a multiplexed sensing system (Figure 1a). The multiplexed nature of TETRIS allows simultaneous sensing of different analytes in the same sample, including concentrations of salts (*via* EIS), H_2_O_2_ and pH, where we chose these analytes due to their relevance as signals in plant physiological processes and stress response (Figure 1b-e).^12^ The sensing module, holding a single sensor (or multiple sensors for multiplexed measurements), was placed into the chamber, consisting of a silicone base and transparent acrylic lid (**Figure 2a**). A reservoir of water was created in the silicone base to ensure that a level of nearly 100% relative humidity was maintained to prevent evaporation of water from the root environment and sensor, and therefore prevent erroneous measurements during the test (see supplementary information **Figure S1**). The sensors were connected to a standard laboratory potentiostat (PalmSens 4 by PalmSens BV, Netherlands) with two 8-channel multiplexers for multiplexed measurements. The potentiostat was connected to a desktop PC for real-time measurements.

**Figure 2.**
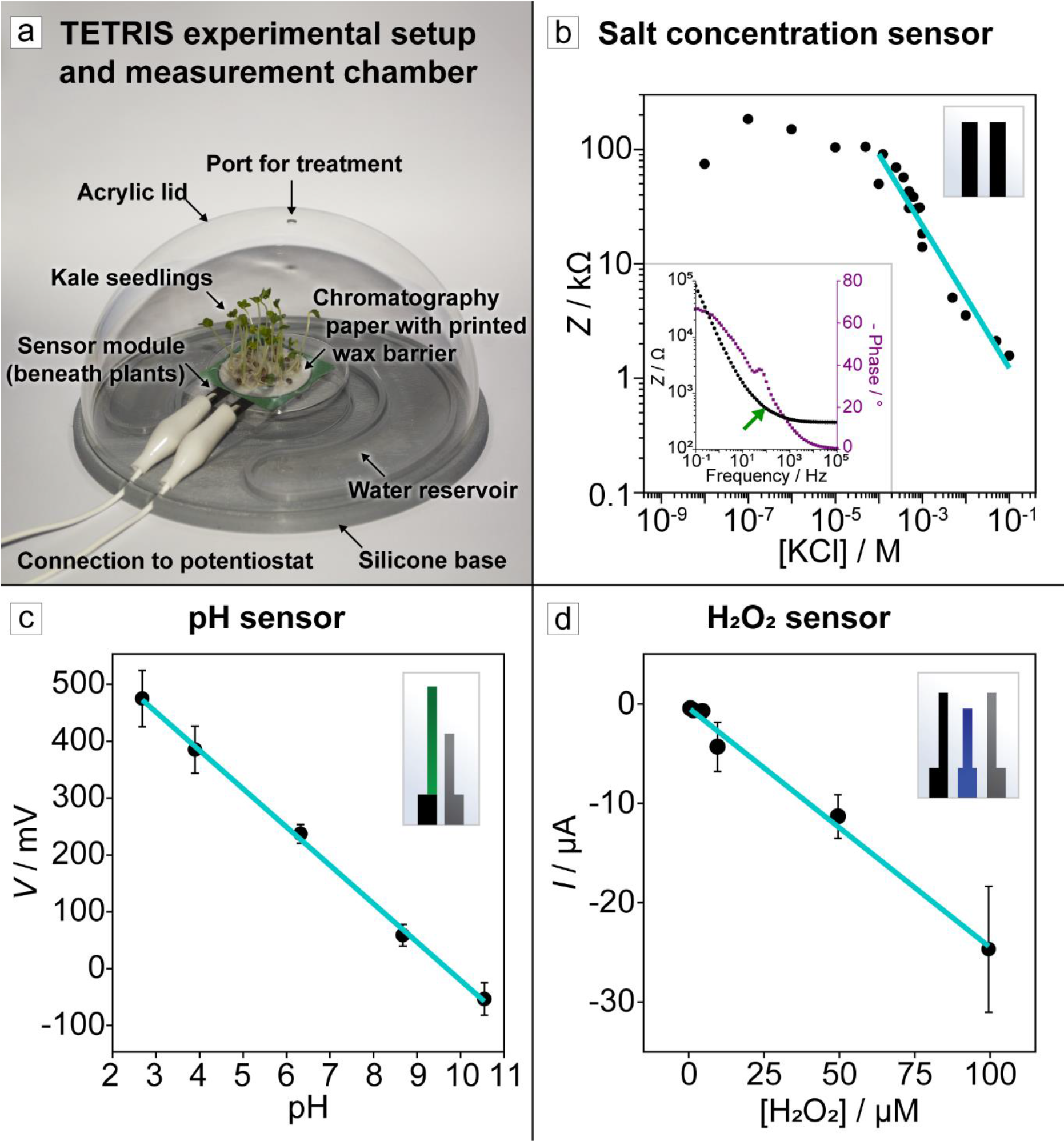
a) Photograph of TETRIS, where single or multiple sensors can be placed under seedlings grown on paper; **b)** Average impedance response of our salt concentration sensor to various concentrations of KCl at 2 kHz. Solid teal line indicates linear fit of linear range of data. Insert shows Bode plot of electrochemical impedance spectrum in 0.1 M KCl, where green arrow shows our chosen frequency of 2 kHz; **c)** Calibration curve of pH sensor in 1 M KCl, error bars show 1 standard deviation (n = 8); **d)** Calibration curve of H2O2 sensor in 1 M KCl, error bars show 1 standard deviation, (100 µM: n = 4; 2 µM: n = 2; 5–50 µM: n = 3). Sensor design displayed in the top right corner of each plot b) – d).

All experiments were performed with the plants grown on discs of chromatography paper (Whatman 1). Seeds were first germinated on wet tissue for two days and transferred onto chromatography paper, where hydrophobic wax barriers were created by a Xerox ColorQube solid wax printer to define the growth and measurement area of the paper disc. The presence of a wax barrier also prevented the analyte solution from reaching the contacts and shorting the measurement circuit. The plants were grown in enclosed boxes with a constant water supply for a set number of days, in batches large enough for each experiment. The paper substrate with the plant seedlings was then placed onto the sensor module to start the measurement.

### Characterization of sensors

We designed three sensors for use in TETRIS: i) an impedance-based non-specific salt concentration sensor, ii) a potentiometric pH sensor and iii) an amperometric H_2_O_2_ sensor (Figure 1b-e, **Figure S2**). Our low-cost (∼$0.02 per sensor) salt concentration sensor consisted of two screen-printed carbon electrodes on polyester transparency sheet. By measuring the impedance of the solution between the carbon electrodes with an excitation signal of 2 kHz, 0.25 V_half wave_ (RMS), we were able to monitor the overall, non-specific salt concentration in the paper substrate. Calibration experiments for the impedance-based sensor were carried out for salts consisting of a range of cations (Ag^+^, Ba^2+^, Ca^2+^, Cd^2+^, Cu^2+^, Gd^3+^, K^+^, La^3+^, Mg^2+^, Na^+^, Ni^2+^, NH_4_ ) and anions (Cl , CO_3_ , PO_4_ , NO_3_ , OH , SO_4_ ) on paper discs. A relationship was found between the logarithm of total electrical impedance (*Z*) and the logarithm of salt concentration (*c*), where impedance decreased with increasing salt concentration.

Calibration curves were found for all the salts by taking measurements from 0.125–1.00 mM (**Table S1**). Extended studies for some salts (Table S1) demonstrate that this log(*c*)-log(*Z*) relationship extends down to 0.1 mM and up to 0.1 M, as shown for KCl in **Figure 2b**. The response time of the sensor after the addition of salt to paper was fast (faster than the frequency of measurement the potentiostat allowed), followed by a lengthier stabilization time before a steady impedance value was reached, likely due to ion diffusion through the paper (Figure S2).^34^ Addition of deionized water did not have any observable effects on the overall impedance of paper that was already wet. Differences in total volume of solution in the paper were found to have only a small effect on the measured impedance, with considerably higher impedance only observed for low volumes (<200 µL), where significant evaporation occurred (Figure S2) within hours (<5 hours until visibly dry) for these smaller volumes. This emphasized the importance for a controlled measurement chamber with high relative humidity (≥80%) to prevent evaporation and therefore erroneous measurements. The silicone base, acrylic lid and water reservoir did slow down evaporation in the paper, although some evaporation continued to occur in the experiments, leading to a slight up-trending drift in impedance over time. We were able to achieve good reproducibility between sensors for KCl concentrations in the range of 0–0.01 M (standard deviation between values for each concentration ≤ 0.083 for log(*Z* / Ω), n = 5 each at five concentrations) as shown in Figure S2.

The pH sensor produced for TETRIS consisted of two screen-printed electrodes. pH was measured with open-circuit potentiometry, which measures the open-circuit voltage between two electrodes, once sensitive to protons in solution. The working electrode (WE) consisted of printed carbon with a layer of polyaniline (PANI) electropolymerized onto the electrode surface. PANI facilitated pH measurement through protonation and deprotonation of nitrogen atoms on the polymer chain with decreasing and increasing pH, respectively, leading to a change in surface charge and, therefore, electrical potential. The reference electrode (RE) was formed of silver/silver-chloride ink (60:40). We found that our pH sensor produced a linear response of -67.5 ± 1.7 mV per pH (r^2^ = 0.973) between pH 2.7 and 10.5 in 1 M KCl bulk solution by addition of NaOH and H_2_SO_4_ (**Figure 2c**), comparable to other printed pH sensors in the literature.^35, 36^ This slightly super-Nernstian response has previously been reported in PANI-based pH sensors, possibly due to surface hydration effects at time of formation of the PANI layer.^35, 37, 38^ Our sensors displayed long-term stability suitable for our use-case (where the sensors were used for up to ∼100 hours), with an average drift of only 0.68 mV hour^-1^, or -0.010 pH hour^-1^ (Figure S2). While calibration experiments were performed in bulk solution to enable the use of a standard pH electrode, we observed the same pH behavior for our sensors using paper disc substrates.

The H_2_O_2_ sensor in TETRIS consisted of three screen-printed electrodes: a Prussian blue-mediated carbon WE, a Ag/AgCl RE and a carbon counter electrode (CE), measured amperometrically at 0 V *vs* the printed pseudo-RE. The calibration experiment was performed in 1 ml of 1 M KCl on the paper discs. We found a sensitivity of -0.24 ± 0.02 µA µM^-1^ between 2 and 100 µM H_2_O_2_ for our sensor with a WE geometric area of 30 mm^2^ (**Figure 2d**). The measurements produced a linear relation between the current reading and concentration which is in agreement with the previous reports in the literature for this system.^39^ We observed larger variations in the signals measured at higher concentrations (standard deviation = 6.3 µA at 100 µM H_2_O_2_, 0.32 µA at 1 µM H_2_O_2_), possibly due to variations in the geometry of the electrodes between each sensor and variation in the diffusion of H_2_O_2_ through the paper. Extracellular H_2_O_2_ concentration due to salt stress, metals or other chemical stimuli can change in the order of 0.1 µM to tens of µM, a range covered partly but not fully by our sensor.^40^ To further increase the sensitivity, we could potentially roughen the electrode surface to increase surface area, deposit nanoparticles of Prussian Blue or fully optimize the appropriate voltage.^41^ Constraints remain in the general setup, however, where diffusion of H_2_O_2_ is lowered by the paper substrate and could provide a limit of detection higher than the sensor would otherwise be capable of.

### Continuous measurement of chemical signals in the plant root environment

We used TETRIS to monitor in real-time the chemical environment in the roots of plants grown on paper. After growing for a certain time, the paper disc with seedlings was removed from the growth box and placed onto the sensing module (Figure 2a). **Figure 3** shows the continuous measurement of impedance, pH and H_2_O_2_ in this setup. **Figure 3a** displays the addition of H_2_O_2_ to the roots of plants, where the increase in signal upon addition is visible. The control (chromatography paper composed of cotton cellulose, with no kale seedlings) showed a larger peak than with kale seedlings upon addition of H_2_O_2_ (30 µL, 500 µM) to the paper substrate. This difference in signal could be attributed to enzymes produced by the roots removing H_2_O_2_, where peroxidases are especially abundant in roots, and the physical presence of the roots slowing down diffusion.^42^ Such a sensing system could be utilized for detecting burst of reactive oxygen species, such as H_2_O_2_, which is released upon detection of pathogens by the plant.^29, 43, 44^

**Figure 3.**
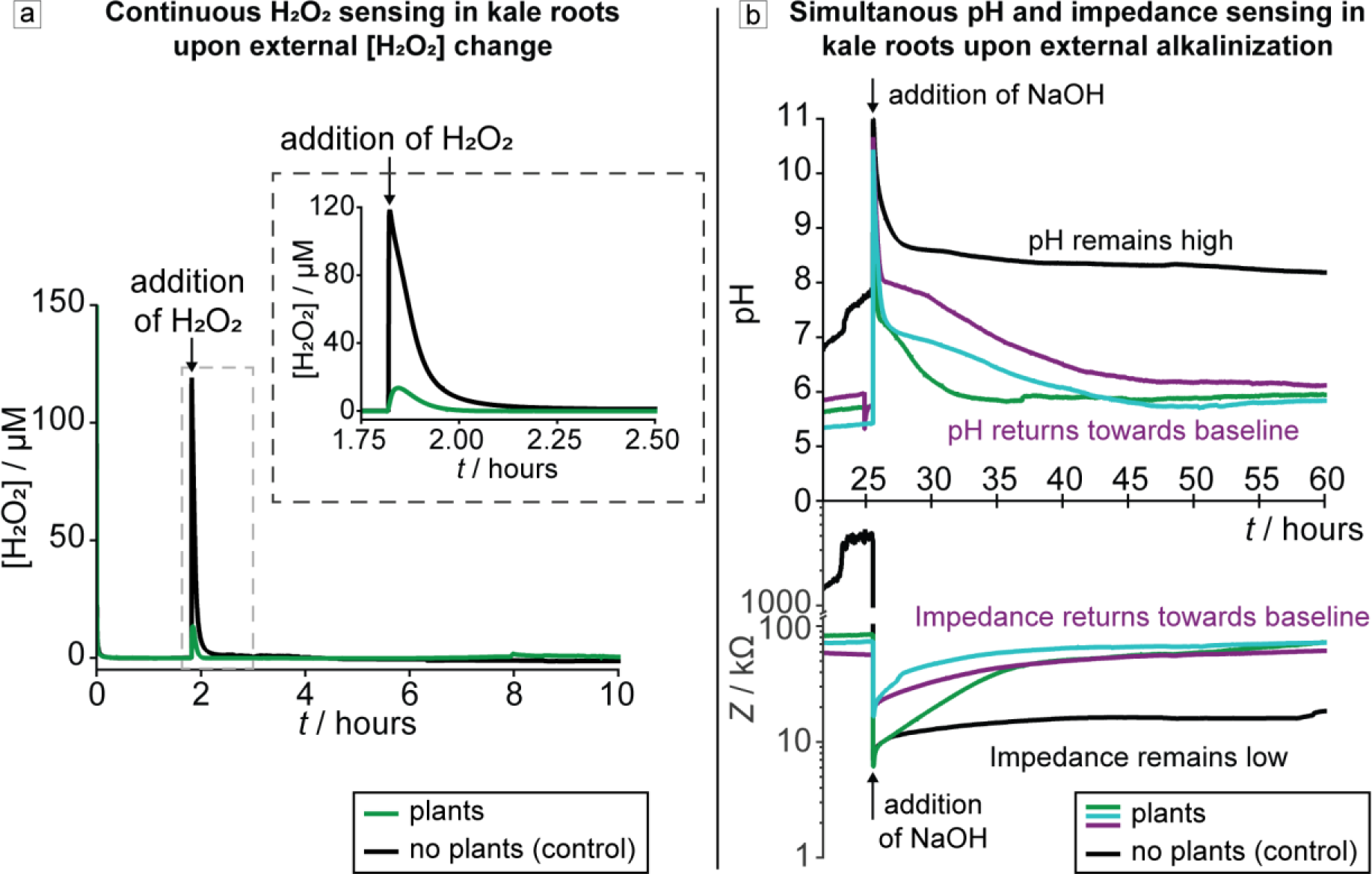
a) Real-time monitoring of H2O2 concentration in the roots of kale seedlings (green signal, 10 plants, 16-days-old) and blank paper control (black signal), with addition of H2O2 (500 µM, 30 µL) into the paper disc (black arrow). Dashed border inset shows magnified plot. **B)** Real-time, simultaneous monitoring of pH (top) and electrical impedance (bottom) of the roots of kale seedlings (green, purple and cyan signals, 30 plants, 9-days-old) and blank paper control (black signal), with addition of NaOH solution into the paper disc (black arrows). Data smoothed in OriginPro using a percentile filter (50^th^ percentile, 30 points of window).

We also simultaneously measured real-time impedance and pH changes, placing the two sensors adjacently, where our aim was to demonstrate the multiplexing capabilities of TETRIS (**Figure 3b**). We modulated the external pH by adding a basic solution of a different pH to observe any time-dependent effects on the pH and impedance of the sample. Initially, the kale samples had a lower impedance and pH than control samples (chromatography paper discs without kale seedlings) due to the presence of nutrients in the seeds and plant root exudates, including ions, amino acids, hormones and other molecules.^45–47^ The pH of the samples was adjusted with NaOH solution (pH 9, 300 µL), performed to simulate saline-alkali stress (previously explored in the literature for studying the effects of damage to roots) and the pH value chosen to match the value of soil considered alkaline.^48, 49^ A rise in pH and a lowering in impedance was initially observed for all samples. Over the course of the following 35 hours, however, we observed in the kale samples a lowering of pH below 6.5 and increase in impedance back towards their respective initial baseline values. Meanwhile, the pH of the control sample remained above pH 8 for at least 30 hours after addition and the impedance remained below 20 kΩ. The increase in impedance observed in the kale samples was attributed to salt uptake into the plant by the roots. Plants have evolved a variety of mechanisms to take up nutrients through the roots, including primary active transport (adenosine triphosphate (ATP) moves nutrients across membranes, going against the nutrient’s gradient), secondary active transport (H^+^-ATPases drives uptake through existing gradients by creating electrical and proton gradients), membrane-bound transport proteins, and passive diffusion.^33^ It is likely that the Na^+^ ions were taken up by the roots of the plants and as charge balance must be maintained, the root cells release H^+^ ions (mainly through H^+^-ATPase) to maintain this balance.^50^ These extruded H^+^ ions may then neutralize OH^-^ ions located at the root surface, reducing the pH.

### The uptake of KNO3 by kale seedlings

Next, we investigated the increase in impedance observed over time in Figure 3b in more detail using TETRIS to understand the uptake dynamics of various inorganic salts by kale seedlings (Video S2). A real-time impedance measurement with added KNO3 can be seen in **Figure 4a**, where the measured impedance after adding salt solution to kale is compared to addition of water to kale and addition of salt to chromatography paper only (composed of cotton cellulose, without seedlings). After an initial period for stabilizing the response of the sensors, addition of 30 µL water onto the paper substrate containing the seedlings did not cause a significant change in impedance, whereas addition of KNO3 (0.1 M, 30 μL) led to a large drop in impedance; a drop of ∼60 kΩ was observed for seedlings and a larger drop of ∼260 kΩ was observed without seedlings due to the higher initial starting impedance.

**Figure 4.**
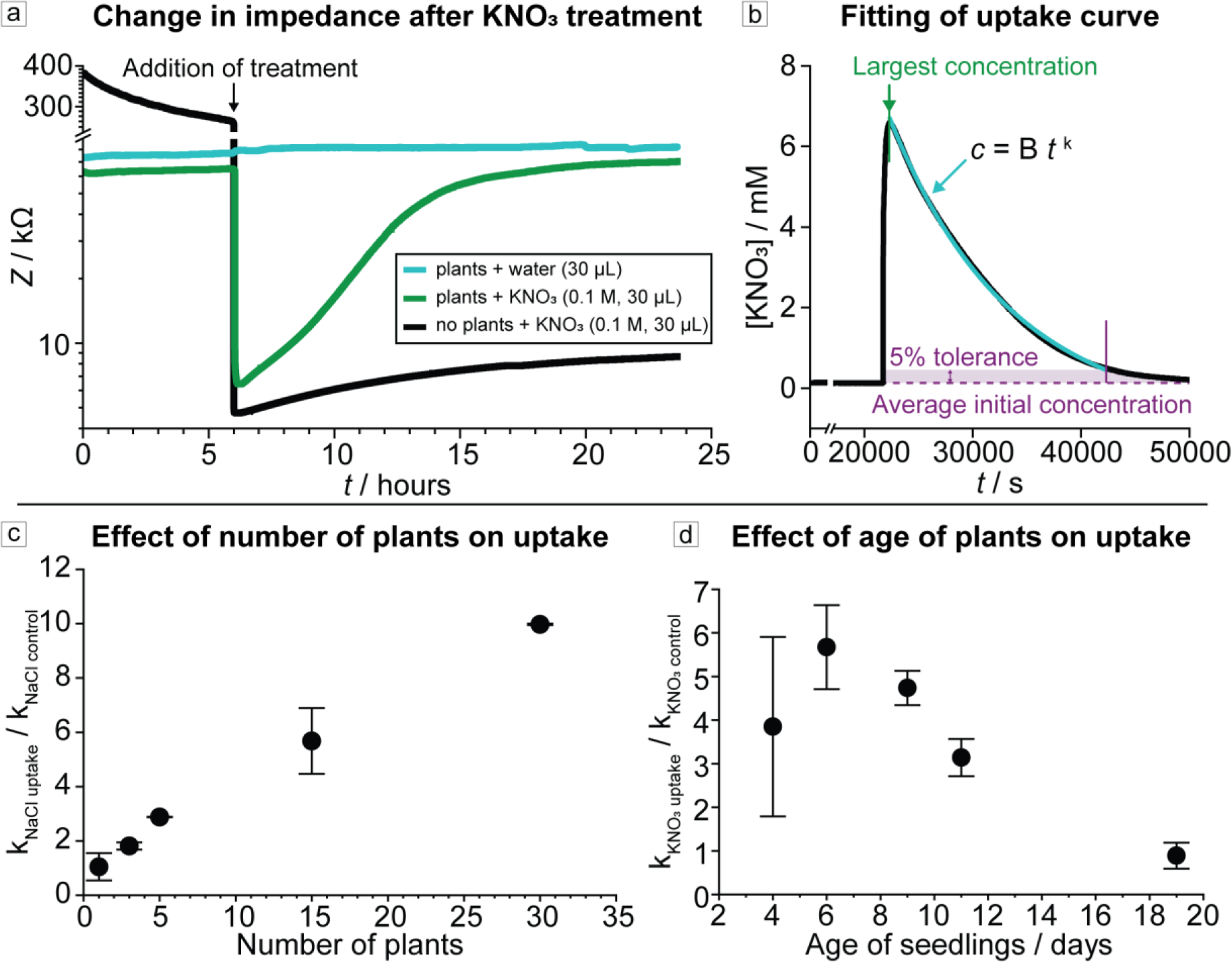
a) Kale seedlings grown on filter paper were placed onto the impedance sensor within TETRIS and the electrical impedance measured in real-time. KNO3 (0.1 M, 30 μL) was then added to the filter paper after a rest period of at least 3 hours (arrow). The impedance slowly increased as ions were taken up by the seedlings and the salt concentration in the paper decreased (green signal). This is compared to empty paper substrate (no plants present), where the impedance remained low (black signal), and plants on paper with only water added instead of salt solution, where the impedance remained high (teal signal). Data smoothed in OriginPro using a percentile filter (50^th^ percentile, 100 points of window); **b)** An exponential curve of the form *c* = B*t*^k^ was plotted between the point of largest salt concentration measured up to where the curve meets the initial baseline concentration, with 5% tolerance. k was found and set as the uptake amount; **c)** Effect of number of 9-days-old kale seedlings on relative amount of uptake of NaCl. Error bars indicate 1 standard deviation (n = 2); **d)** Effect of age of kale (thirty plants) on relative amount of KNO3 uptake. Error bars indicate 1 standard deviation (n = 3).

The impedance remained steadily low in the blank filter paper without any seedlings, but in the sample with kale seedlings the impedance increased over time, reaching the initial impedance level of the sample, suggesting uptake of KNO3.

We first produced calibration curves for the salt concentration sensors which correlate the impedance readings to salt concentrations (Table S1). To then determine the change in salt concentration due to uptake, concentration (*c*) was plotted against time (*t*) and a logarithmic curve was fitted of the form *c* = B*t*^k^ (where B is a constant) to find the exponential term kuptake, the rate of uptake of the salt (**Figure 4b**). As evaporation of water produced a small increase in impedance over time (∼0.1 kΩ hour^-1^, see black signal in Figure 4a), to make a true comparison between different salts, kuptake was normalized to the increase in impedance due to evaporation (kcontrol). kcontrol was found in the same way as kuptake, but where no plants were grown on the paper.

We characterized the rate of uptake of NaCl (addition of 0.1 M, 30 μL) *vs* number of kale seedlings grown on the paper substrate as shown in **Figure 4c**. With the increasing number of plants, the collectively measured rate of uptake also increased in a linear fashion, likely due to increased root surface area and higher overall collective rates of transport. Although this experiment produced results that were expected, strong evidence was generated, confirming the functionality of TETRIS for measuring rates of uptake for dissolved ions.

Next, we studied the relationship between the rate of uptake of salts and the age of plants used in the experiment **(Figure 4d)**. We observed that the rate of uptake of salt increased with the age of plants, up to plants aged six days (their age from germination until the start of the experiment), albeit with large variation that was likely due to the variation in root growth of the young plants. For plants aged between six days and 19 days, the rate of uptake of salt decreased with the age of the plants used in the experiment. At 19 days old, the rates of uptake with plants were not significantly different from the paper control without plants. This was a surprising finding; although the older plants were visibly larger than younger seedlings, the rates of uptake were lower. Although we do not know exactly why, we speculate that because we do not provide any nutrients to the seedlings, other than deionized water during growth, the seedlings may not be developing as normal. This notion may also be supported by the observation that after about two weeks of growth on paper, some wilting, discoloration and decay were generally evident in the plants. This plant degradation could have occurred due to microscopic cell death or lignification of the root tissue.^51^ Furthermore, unlike soil, paper is not a three-dimensional substrate and hence it does not provide the three-dimensional mechanical matrix that presumably allows support for proper root development or plant growth.^52^ In any case, the underlying biology of this observation will need to be studied in future experiments.

### Comparing the rates of uptake of nutrients, heavy metals and sodium salts

To characterize the rates of uptake of different ionic species, kale seedlings were treated with salts containing a range of cations and anions to observe the effect. The change in salt concentration in paper discs with thirty kale seedlings was determined using TETRIS for each salt and compared to the recorded change in concentration for plain paper discs with no plants. Plants aged nine days were selected due to their high rates of uptake without large deviation between samples (Figure 4d).

Normalized rates of uptake of salt were more reliable than individual rates as processes such as evaporation caused a slight increase in impedance over time, as outlined before. A clear difference in normalized rate of uptake was observed between chemical species (**Figure 5a**). Salts considered to be nutritious for plants (macronutrients: K^+^, NH4 ; secondary nutrients: Ca , Mg ) or those with smaller cations (Na^+^) exhibited higher rates of uptake than those salts with heavy metal (HM) cations (Ag^+^, Ba^2+^, Cd^2+^, Cu^2+^, La^3+^, Ni^2+^). By separating the ions into those categories (macronutrients, secondary nutrients, sodium, HMs), we found statistical significance in the average rate of uptake according to ion category by one-way ANOVA (f(3,111) = 10.9353, p < 0.0001). A Tukey post-hoc test revealed significant pairwise differences between some of the categories, including between both macro and secondary nutrients and HMs, and the confidence intervals and means are displayed in **Figure 5b**.

**Figure 5.**
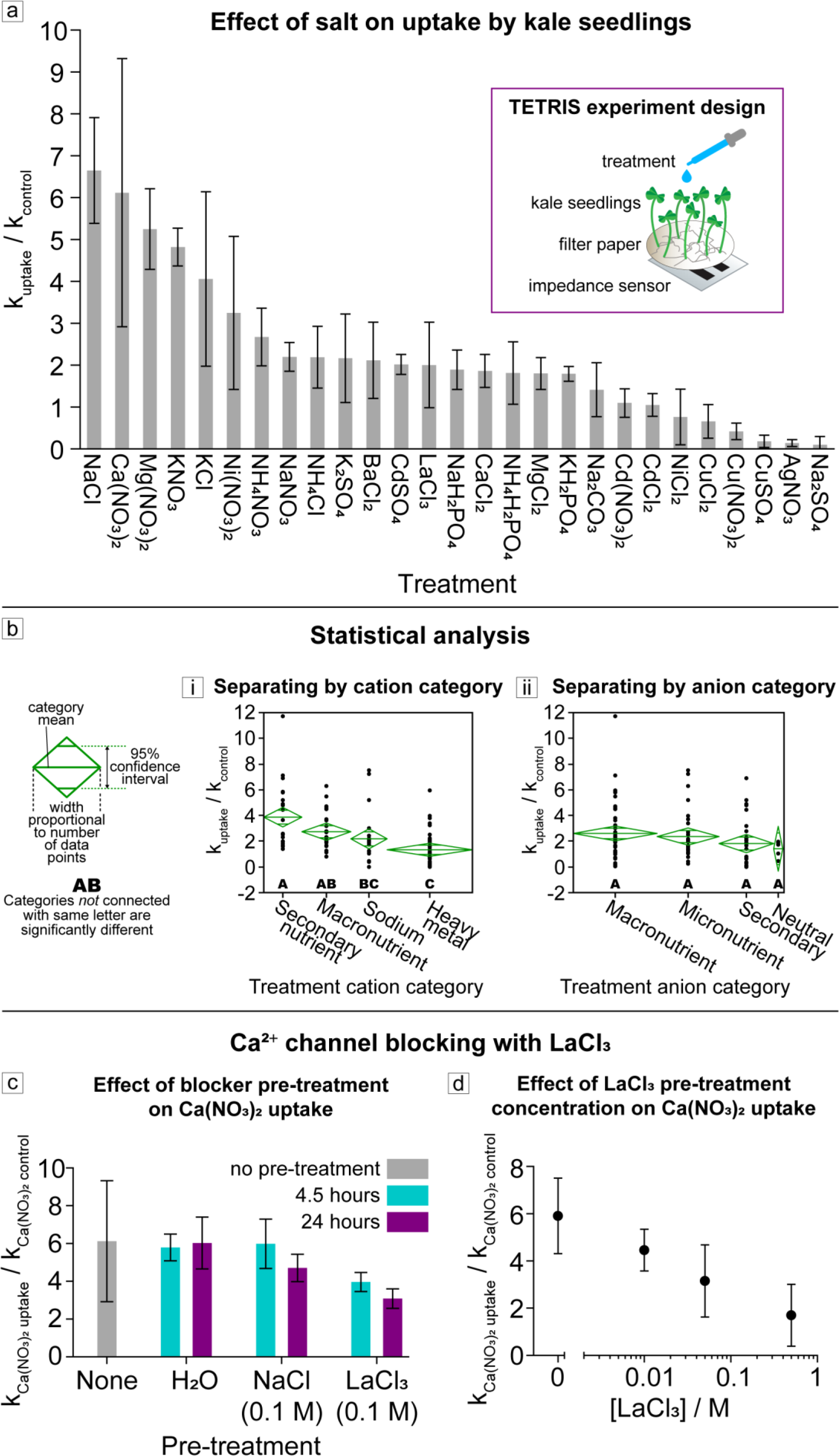
a) Rate of uptake of salt from paper disc with thirty kale seedlings (9-days-old), relative to control without plants, using TETRIS. (Values at or below 1 are considered to have a net zero or negative ion uptake). Error bars indicate 1 standard deviation (CdCl2, KCl, NaCl: n = 3; CaCl2, Ca(NO3)2: n = 6; LaCl3: n = 7; all others: n = 4). **b)** Statistical analysis showed significant differences between some cation classes (i), but not between anion classes (ii), where confidence diamonds show mean uptake, 95% confidence interval and number of data points, and letters show categories with or without significant difference. **c)** Pre-treatment of thirty kale seedlings (9-days-old) with deionized water, NaCl (0.1 M) or LaCl3 (0.1 M) for 4.5 or 24 hours and their effect on the relative change in concentration of Ca(NO3)2. Error bars indicate 1 standard deviation. (None: n = 6; LaCl2 24 hours: n = 4; all others: n = 3). **d)** Pre-treatment of thirty kale seedlings (9-days-old) with varying concentration of LaCl3 for 4 hours and the effect on the relative change in concentration of Ca(NO3)2. Error bars indicate 1 standard deviation (n = 2).

While secondary nutrients had a greater average rate of uptake than macronutrients, this difference was not found to be statistically significant.

The differences in rate of uptake can be explained through the different methods of uptake and transportation of ions in plants. K^+^ is a macronutrient for plants (nutrients that are required and consumed in large quantities). K^+^ is taken up by multiple mechanisms, grouped into seven families of K^+^-permeable cation transporters, although not all of these transporters are exclusive to K^+^.^53^ Na^+^, a toxic ion for plants, competes with K^+^ uptake through high-affinity potassium transporters (HKTs) and nonselective cation channels (NSCCs) due to their similar chemical properties and ion sizes, explaining the high rate of uptake observed of Na^+^ salts.^54^ NH4 is a common source of nitrogen for plants (alongside NO3 ); high rates of uptake we observed could be attributed to NH4 transport systems in the root plasma membrane, despite NH4^+^ concentrations rarely naturally exceeding 50 μM in soil.^55^ Ca^2+^ is another important nutrient for plants, with multiple transport mechanisms, including Ca^2+^-permeable ion channels, Ca^2+^-ATPases and Ca^2+^/H^+^ exchangers.^56^ Ca(NO3)2 showed a relatively high rate of uptake, although CaCl2 showed a much slower uptake. This is potentially due to Cl^-^, considered a micronutrient, being taken up less than the macronutrient NO3 . A similar relationship was also observed with salts of Mg^2+^, where Mg(NO3)_2_ showed a relatively high rate of uptake compared to MgCl2. Like Ca^2+^, Mg^2+^ is a key nutrient, with specific transporters for uptake via the roots, and so we observed similar rates of uptake to Ca^2+^.^57^

Many HM ions are encountered by plants through human-derived industrial and agricultural activities. Excessive HM uptake can be detrimental to the health of the plant through oxidative stress, damage to cell structures and substitution of nutrients at cation exchange sites.^58^ Consumption of crops containing toxic HMs by humans can lead to health problems through accumulation of HMs in the body, indicating the importance of investigating plant uptake of HM ions. Furthermore, specific plants are often used to remove HMs from contaminated soil through “phytoremediation”, and therefore plants that exhibit high rates of uptake of HMs are just as important as those that do not.^59^

The rate of uptake would be expected to be low for most HMs due to a lack of transporters for non-essential HM ions, as was generally observed in our experiments (Figure 5). HM ions, depending on their species, may be considered micronutrients (for example, Cu^2+^ is necessary for photosynthesis and Ni^2+^ activates urease activity) or potentially toxic pollutants, or sometimes either depending on the concentration. Essential HM ions may have specific transporters, but non-specific diffusion-based uptake can occur for most HM ions, depending on the surrounding concentration. Ag(NO3)2 was found to have a negligent amount of uptake, potentially due to its reduction to metallic silver, silver oxide or AgCl, as suggested by the appearance of a gray discoloration forming on the roots of the plants (**Figure S3**).^60^ Na2SO4 also displayed negligible uptake, a surprising observation as Na^+^ and SO4 (as a source of S) both typically displayed high rates of uptake. A previous study found that transport rates of Na^+^ are higher in treatments containing Cl^-^ than SO4 , corroborating the lower rate of uptake for Na2SO4 compared to that for NaCl, but such a low rate for Na2SO4 was unexpected.^61, 62^ Cd^2+^ salts generally also had an observed low rate of uptake. It has previously been reported that cadmium is absorbed on the surface of the root but not beyond, suggesting that a minimal amount of cadmium is taken up by plants, in agreement with our observations.^60^

Whereas separating the salts into their cation’s relationship to plants revealed statistical significance in rate of uptake, we did not find a similar significance when separating the salts by their anions. All the anions used here are considered either macronutrients (NO3 , PO4 ), secondary nutrients (SO4 ), micronutrients (Cl^-^ ) or neutral (CO3 ). None of these anions are especially dangerous to plants (not even Cl^-^, which recently has even had a re-evaluation as a possible macronutrient) and would only be considered toxic in large concentrations (4–35 mg g^-1^ dry weight).^61, 63–65^ Anions are also typically absorbed in excess compared to cations, thus are unlikely to be the limiting factor in salt uptake, and so it is perhaps not surprising that we did not observe significant differences in uptake due to anion.^50^

### Ca^2+^ channel blocking by LaCl3

The blocking of cation channels is of great interest for scientists looking at stress tolerances and reactions in plants. For example, internal processes (such as the result of wound response *via* effects of inducers, like systemin, on Ca^2+^ signaling) and external processes (such as salt or nutrient stress *via* Na^+^ and nutrient uptake by root cells) can both be probed by using ion channel blockers.^6,66^ Certain HM ions can block the ion channels of smaller ions; for example, BaCl2 has been shown by radiotracer experiments to decrease sodium uptake, and Ca^2+^ channels can be blocked by LaCl3.

We investigated the LaCl3 blocking effect using TETRIS by pre-treating plants with a blocker or a control prior to monitoring salt uptake. The paper and seedlings were then washed in deionized water to remove excess blocker and the plants transferred onto fresh, pre-wetted paper discs. The impedance measurement was then initiated and, after a rest period of 24 hours, a 30 μL solution of 0.1 M Ca(NO3)_2_ was added. Initially, the effect of pre-treating plants with LaCl3 (0.1 M), a known Ca^2+^ channel blocker, was compared to the effect of pre-treating plants with NaCl (0.1 M), a salt not considered to block Ca^2+^ channels, and deionized water (**Figure 5c**).^66^ Compared to the normalized rate of uptake for Ca(NO3)2 without treatment (6.1 normalized uptake), we found similar rates of Ca(NO3)2 uptake when pre-treating the plants for 4.5 hours with deionized water (5.8) or NaCl (6.0). Pre-treating plants with LaCl3, however, led to a reduced rate of uptake for Ca(NO3)2 of around 4.0 (about a third reduction in uptake compared to the control). Increasing this pre-treatment time from

4.5 to 24 hours, we found that a DI water treatment produced a similarly high normalized rate of uptake for Ca(NO3)2 (6.0), but the addition of NaCl or LaCl3 did not. Treatment with NaCl led to a reduced rate of uptake of around 4.7 (a reduction of about a fifth), suggesting that NaCl may provide some blocking effect with longer treatment. It is also possible that excess NaCl remains on the surface of the roots between transfer to new paper disc. In any case, the blocking effect of LaCl3 on was still far more pronounced; increasing LaCl3 pre-treatment time to 24 hours reduced the normalized rate of uptake of Ca(NO3)2 to 3.1, around half the uptake of the control.

Using TETRIS, we also investigated the effect of the concentration of LaCl3 on blocking Ca^2+^ channels. We prepared solutions of LaCl3 with four different concentrations ranging from 0–0.5 M. Next, the kale seedlings were pre-treated for four hours with the LaCl3 solutions prepared before measuring the rate of uptake for Ca(NO3)2 using printed impedance sensors. We observed a marked decrease in the rate of uptake for Ca(NO3)2 with the increasing concentrations of LaCl3 (**Figure 5d**) in a log-linear fashion. Although we did not go beyond 0.5 M, further increases in the concentration of the blocking agent LaCl3 or duration of pre-treatment is likely to improve blocking of Ca^2+^ channels, though clearly to a lesser extent.

The reduction of uptake of Ca(NO3)2 with LaCl3, but not complete prevention of uptake, suggests that cellular uptake may not be the only possible route of ion uptake. The seedlings also potentially absorb water, containing the ions from Ca(NO3)2, into xylem vessels. As the La^3+^ only blocks ion channels, uptake into the xylem from the roots *via* diffusion is still unhindered, and so some uptake of Ca(NO3)2 may have still occurred.^33^

With these results, we have shown the potential application of printed impedance sensors for studying continuous-time ion transport in plants, demonstrating the utility of printed impedance sensors in applied and fundamental plant research.

### Predicting rate of uptake of salts with machine learning

We developed a supervised machine learning model for TETRIS for predicting the rates of uptake for salts using only the physicochemical properties and the biological relevance of the salts as inputs to the model to accelerate plant phenotyping. This approach may be most useful in identifying new regulators of ion uptake mechanisms and of cellular stress responses. This would subsequently lead to the development of new plant varieties with higher yields and higher tolerance to environmental fluctuations caused by climate change or soil salinization. Machine learning techniques (including neural networks and predictive clustering trees) have been used previously to estimate nutrient uptake of herbage in fields and water uptake in roots, but a modelling approach has not been used to monitor and predict uptake of salt on the individual plant scale.^67, 68^

We used the eXtreme Gradient Boosting (XGBoost) algorithm for the development of the supervised machine learning model to predict the normalized rate of uptake (kuptake / kcontrol). In simple terms, XGBoost is a scalable machine learning system for tree boosting – a method of combining many weak learners (or “trees”) into a strong classifier. The XGBoost method aims to find a relationship between the input X = {x1, x2, …, xN} and the output Y = {y1, y2, …, yN}.^69, 70^ We built five models to predict kuptake / kcontrol within different ranges, from a simple two-range “uptake/no uptake” model to predicting kuptake / kcontrol with increasing resolution. We trained the model using the rates of uptake for the salts measured and physicochemical information on the individual ions in the salts, such as their chemical groups, relative masses, ion charges and the biological relevance of the salt to the plant (such as if the ions were deemed to be nutrients) – see Methods section for the full list of features used in the model.^71^ The experimental dataset was split into five folds for cross-validation: in each validation, four folds (∼80% of the data) were used for training and one fold, unseen data (∼20% of the data), for testing, and cross-validation carried out by changing which fold was used for testing. The classification models were evaluated using the *F*1 score (averaged over the different iterations of the cross-validation) – a metric for the accuracy of the model combining precision and recall, defined as: *F*1 = (2 · precision · recall) / (precision + recall) We found that by using this machine learning approach we were able to predict salt uptake with data from less than 100 experiments in the training datasets with reasonably high accuracy.

Confusion matrices with five different range configurations can be seen in **Figure 6a**. The two-range model, aiming to predict whether uptake did or did not occur (kuptake / kcontrol being >1 or ≤1 respectively), had an *F*1 score of 0.949, where “1” would be the perfect score for classification.

**Figure 6.**
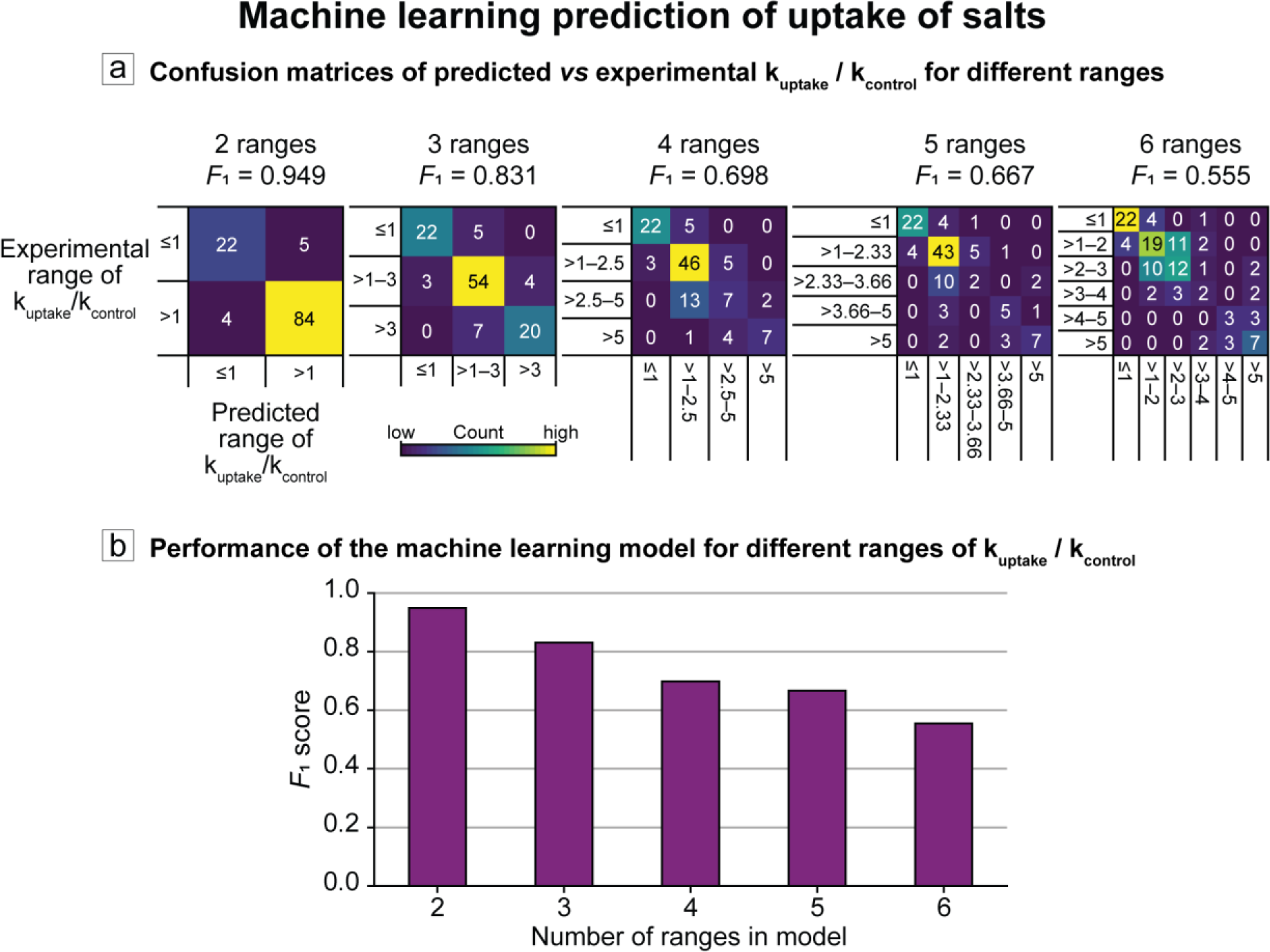
Using machine learning algorithms to predict normalized rate of uptake (kuptake / kcontrol). **a)** Confusion matrices comparing predicted and experimentally obtained ranges of (kuptake / kcontrol). **b)** *F*1 score (a metric of accuracy combing precision and recall) decreases with increasing number of ranges in the machine learning model.

Expectedly, increasing the number of ranges from two to six led to a decrease in *F*1 score (**Figure 6b**) with six ranges producing the lowest score of 0.555.

## 4. Conclusions

TETRIS is a low-cost platform for accelerated phenotyping for plants. As electrochemical root measurements are not utilized with living whole plants, TETRIS provides an important stepping-stone towards the measurement of physiological changes in whole plants, which often have different behavior to individual plant organs or tissue samples. The roots are especially neglected in research, despite being the entry point for uptake of beneficial substances (water, nutrients) or harmful chemicals (sodium salts, heavy metals, extreme pH). By measuring in real-time, we not only enable information-rich observations into time-dependent chemical signals, but we unlock the potential for discovering new biology.

The entire TETRIS platform can be manufactured using tools, such as 3D, wax and screen printers, that are accessible to most academic laboratories. The total cost of materials to produce a single TETRIS system is US $5, excluding the multichannel potentiostat (which can be replaced with a lower cost commercial or a homemade alternative). Although we only performed experiments with kale, the experimental setup was intended as a non-species-specific platform and is compatible with any seedlings that grow on paper or agar gel.

TETRIS can be used to examine the effects of environmental changes or stresses. For example, due to the boxed design, the internal atmosphere could be altered to increase or decrease the concentration of O2 or CO2, gases that are key for plant processes such as photosynthesis and respiration.^27^

The TETRIS platform has the following five disadvantages: i) The electrochemical sensors must remain wet to function and are susceptible to drift if the water reservoir is depleted; ii) The disposable sensors are printed on substrates made of polyester, which are not biodegradable and hence not environmentally friendly. The sensors, however, can be printed directly on paper to overcome this issue;^39, 72–76^ iii) In its current form, the TETRIS platform is not compatible with larger, mature plants, however, because of its scalable characteristic, the platform can be adapted to accommodate other model plants such as *Nicotiana benthamiana*; iv) Measurement in soil may be less feasible due the uneven geometry of soil, however, careful experimental design (such as carefully spreading of soil across the electrodes) may enable measurements in soil; v) The relatively small number of experiments in the dataset (115 uptake experiments) limits the scalability of the machine learning models and has a higher potential for distortion due to outliers. Despite the current limitations, the machine learning approach we have developed could be used for the prediction of rates of uptake of various salts, such as those not previously tested.

Although, in this work, we mostly studied the rates of uptake of ions, with the existing TETRIS setup consisting of printed pH, EIS and H2O2 sensors, biochemical signaling and feedback loops associated with, but not limited to, plant hormones, pesticides, pathogenic or beneficial microorganisms can be studied.^12^ Because TETRIS is modular and customizable, the number of printed chemical sensors integrated into the system can be expanded to include sensors specific to certain ions or molecules.^35^ Furthermore, TETRIS can be used as an artificial rhizosphere to study root microbiome to investigate plant-microbe interactions, and potentially allow creations of digital twins of these complex systems.^77, 78^ In the future, TETRIS can provide a standardized, quantifiable and (with the help of artificial intelligence) autonomous solution to accelerate plant phenotyping. TETRIS has the potential to overcome the urgent “bottleneck” in high-throughput screening in producing high yielding plant.^79, 80^

## 5. Materials and methods

### Plant growth

A hydrophobic wax border was printed (Xerox Colorqube 8580) onto chromatography paper (Whatman, grade 1 chromatography paper, 0.18 mm thickness) and heat transferred (Vevor HP230B) to define a circular area (radius 17.5 mm, area 962 mm^2^). *Brassica oleracea acephala* (“dwarf green curled kale”, Mr Fothergill’s) seeds were stirred in 3% hydrogen peroxide (Millipore) for 5 minutes, rinsed in deionized water, and germinated on damp tissue paper for 2 days prior to transfer onto the wax-printed chromatography paper discs. The paper discs with seedlings were placed onto platforms inside a propagator box with transparent lid and the paper was constantly supplied deionized water from a reservoir through paper strips.

### General experimental setup

During measurements, the sensor was adhered to the base of a petri dish (55 mm diameter), placed into a measurement chamber consisting of a silicone base with a reservoir of water, and connected to a potentiostat with crocodile clips. A transparent colorless acrylic lid was placed over the experimental set up and the impedance of the sensor with the sample placed on top measured. All electrochemical experiments were performed using a PalmSens4 potentiostat and MUX8-R2 multiplexers.

### Humidity and temperature of measurement chamber

The relative humidity and temperature were measured inside and outside of the measurement chambers using the Adafruit BME280 (Bosch) I2C or SPI Temperature Humidity Pressure Sensor connected to an Arduino Due and logged using PuTTY.

### Impedance sensor fabrication and experimental setup

Carbon electrodes (Sun Chemical C2130925D1 conductive carbon ink (80 wt%), Gwent Group S60118D3 diluent (20 wt%)) were screen-printed onto polyester transparency sheet (Office Depot).

The sensor design consisted of two identical electrodes (30 mm × 6.25 mm) separated by 5 mm (**Figure 2**). Impedance measurements (amplitude 0.25 Vhalf wave (RMS), frequency 2 kHz, 0 V d.c.) were performed.

### Impedance sensor characterization

*Calibration for each salt*: The impedance was measured between 0.125–1.00 mM (1 mL, n = 8) for the following compounds on paper discs: BaCl2, CaCl2, Ca(NO3)2, CdCl2, Cd(NO3)2, CdSO4, Cu(NO3)2, CuSO4, GdCl3, K2SO4, LaCl3, Mg(NO3)2, MgCl2, Na2CO3, Na2SO4, NaOH, Ni(NO3)2, NiCl2. An extended range (0.01 μM–0.1 M , n ≥ 15) was measured for the following salts: AgNO3, CuCl2, KCl, KH2PO4, KNO3, NaCl, NaH2PO4, NaNO3, NH4Cl, NH4H2PO4, NH4NO3. Impedance was measured for at least 2 hours and the average of the impedance response recorded. log10(*Z*) was plotted against log10(*c*) and the linear fit found for each salt.

*Response time*: The impedance of a paper disc with 270 μL deionized water was measured. After 60 s, 30 μL KCl (0.1 M) was added.

*Effect of volume*: The impedance of a paper disc with 125–1000 μL KCl (0.1 M) was measured for at least 3000 s.

*Electrochemical Impedance Spectroscopy:* Experiments were run at 0 V DC, 0.25 V AC. For EIS with kale seedlings, the disc with seedlings was placed on the sensor with no added deionized water. For EIS in 0.1 M KCl, the sensor was placed in a solution of 10 ml volume.

### pH sensor fabrication, experimental setup and characterization

Carbon (Sun Chemical C2130925D1 conductive carbon ink (80 wt%), Gwent Group S60118D3 diluent (20 wt%)) and silver/silver chloride (Sun Chemical C2130809D5 (95 wt%), Gwent Group diluent S60530D5 (5 wt%)) electrodes were screen-printed onto polyester transparency sheet (Office Depot). Polyaniline (PANI) was electropolymerized (1 V DC, 100 minutes) from a solution of aniline (0.1 M) and oxalic acid (0.3 M) onto the carbon electrode to form the working electrode (sensing area 2 mm × 40 mm). The silver/silver chloride electrode was the reference electrode (sensing area 2 mm × 18 mm). Open circuit potentiometric measurements performed. Characterization experiments were performed in 1 M KCl in bulk solution (not on paper). The pH was altered by addition of H2SO4 and NaOH. pH was measured using a Hanna Instruments HI 5521 pH meter.

### H2O2 sensor fabrication, experimental setup and characterization

Prussian blue-mediated carbon (Sun Chemical C2070424P2 (80 wt%), Gwent Group S60118D3 diluent (20 wt%)), carbon (Sun Chemical C2130925D1 (80 wt%), Gwent Group S60118D3 diluent (20 wt%)) and silver/silver chloride (Sun Chemical C2130809D5 (95 wt%), Gwent Group diluent S60530D5 (5 wt%)) electrodes were screen-printed onto polyester transparency sheet (Office Depot). The Prussian blue-mediated carbon WE (sensing area 2 mm × 15 mm) was placed between the carbon CE and Ag/AgCl RE (sensing area 2mm × 18 mm each) with a gap of 1 mm. Amperometric measurements were performed at 0 V. Characterization experiments were performed in KCl (1 M, 1 ml) on paper.

### H2O2 plant experiments

A paper disc with 10 kale seedlings was removed from growth chamber 16 days post-germination and placed onto the H2O2 sensor in the measurement chamber. KCl (2.2 M, 136.4 μL) was added to the paper. Current was measured over time at 0 V relative to the printed RE and H2O2 (500 μM, 30 μL, in 1 M KCl) was added to the paper after at least 1 hour.

### Combined pH and impedance plant experiments

A paper disc with 30 seedlings was removed from growth chamber 9 days post-germination and placed onto the sensor module in the measurement chamber, where the pH and impedance sensors were placed at least 10 mm apart. 100 μL deionized water added to the paper. Impedance and pH were measured over time and the pH of the paper altered by addition of NaOH (pH 9, 300 µL).

### Salt uptake experiments

A disc with seedlings was removed from growth chamber after a set number of days and placed onto the sensor in the measurement chamber and 100 μL deionized water added to the paper. After measuring impedance for at least 3 hours, 30 μL salt solution was added to the paper through a port in the lid. To calculate the rate of uptake, impedance was converted to salt concentration by the obtained calibration curves. An initial baseline concentration was found by taking the average concentration of the first 2 hours after measurement began. The time where the concentration was highest after addition of salt was found and a logarithmic curve (of the form *c* = B*t*^k^) fitted from this time. Where the curve would meet the initial baseline (with a 5% tolerance), the constant k was found of the curve up to that timepoint. Where the line did not reach the initial baseline, a curve was plotted until the end of the experiment and the constant k found. Where the concentration did not decrease after addition of salt (and therefore, there was no maximum concentration), the uptake was set to zero. The constant k found for each experiment was then divided by the average k found for the control (no plants) to find the normalized rate of uptake.

### Ion channel blocker experiments

The paper discs with seedlings were removed from the water reservoir after 7 days. 30 μL blocker and 100 μL deionized water were added to the paper. After 4 hours, the discs with seedlings were washed in deionized water and the seedlings transferred to new pre-wetted discs of paper in the measurement chamber. 200 μL deionized water was added to the paper. After measuring impedance for at least 24 hours, 30 μL salt solution was added to the paper through a port in the lid.

### Prediction of salt uptake with machine learning

XGBoost was used to predict the rate of uptake of salts on the following input variables: 1) cation, 2) anion, 3) cation classification (heavy metal, primary macronutrient, secondary macronutrient, micronutrient, sodium), 4) cation wider chemical classification (transition metal, s-block, lanthanum, none), 5) cation chemical group, 6) cation chemical period, 7) cation charge, 8) number of anions, 9) anion charge, 10) anion classification (primary macronutrient, secondary macronutrient, micronutrient, neutral), 11) cation relative mass, 12) anion relative mass, 13) total salt mass, 14) Salt solubility at 25 °C. The categorical features (1-10) were encoded as a one-hot numeric array. For all models, exhaustive searches were performed to identify optimal hyperparameters used, by a 5-fold cross-validated grid-search over a parameter grid. The hyperparameters used for the classification models were as follows: 6 classes (objective: “multi:softprob”, booster: “gbtree”, eval_metric: “merror”, eta: 0.1, max_depth: 3, subsample: 0.7, colsample_bytree: 0.6); 5 classes (objective: “multi:softprob”, booster: “gbtree”, eval_metric: “merror”, eta: 0.1, max_depth: 7, subsample: 0.9, colsample_bytree: 0.6); 4 classes (objective: “multi:softprob”, booster: “gbtree”, eval_metric: “merror”, eta: 0.1, max_depth: 3, subsample: 0.6, colsample_bytree: 0.4); 3 classes (objective: “multi:softprob”, booster: “gbtree”, eval_metric: “merror”, eta: 0.15, max_depth: 3, subsample: 0.7, colsample_bytree: 0.1); 2 classes (objective: “binary:logistic”, booster: “gbtree”, eval_metric: “logloss”, eta: 0.15, max_depth: 5, subsample: 0.6, colsample_bytree: 0.6).

## Acknowledgments

The authors would like to thank the department of Bioengineering at Imperial College London. F.G. and P.C. thank the Imperial College Centre for Processable Electronics (CPE). F.G., P.C. and A.C. thank EPSRC (EP/L016702/1) and BBSRC DTP (Reference: 2177734). F.G. and L.G.-M. thank Bill and Melinda Gates Foundation (Grand Challenges Explorations scheme under grant number: OPP1212574 and Investment ID INV-038695) for their generous support. F.G. thanks the US Army (U.S. Army Foreign Technology (and Science) Assessment Support program under grant number: W911QY-20-R-0022). L.G-M. acknowledges European Union’s Horizon 2020 research and innovation program under the Marie Sklodowska-Curie grant agreement No 101025390. T.B. acknowledges BBSRC (BB/T006102/1) for their support. S.O. acknowledges the Imperial President’s Ph.D. Scholarship. Y. C. would like to thank the Turkish Ministry of Education and EPSRC IAA. F.G. also acknowledges Agri Futures Lab. T.A. would like to thank Innovate UK (grant reference: 10004425).

## Author Contributions

P.C., L.G-M. and F.G. conceived the structure of the manuscript. P.C. lead the writing of the manuscript and experimental work. Y.C., A.C., T.A., S.O., and A.N. contributed to experimental work. L.G-M., F.G., T.B., A.C. and A.N. contributed to the writing of the manuscript. All authors reviewed and agreed on the manuscript before submission.

