## Supporting Information for "Time-Resolved Chemical Phenotyping of Whole Plant Roots with Printed Electrochemical Sensors and Machine Learning"

### for

### Supporting Information

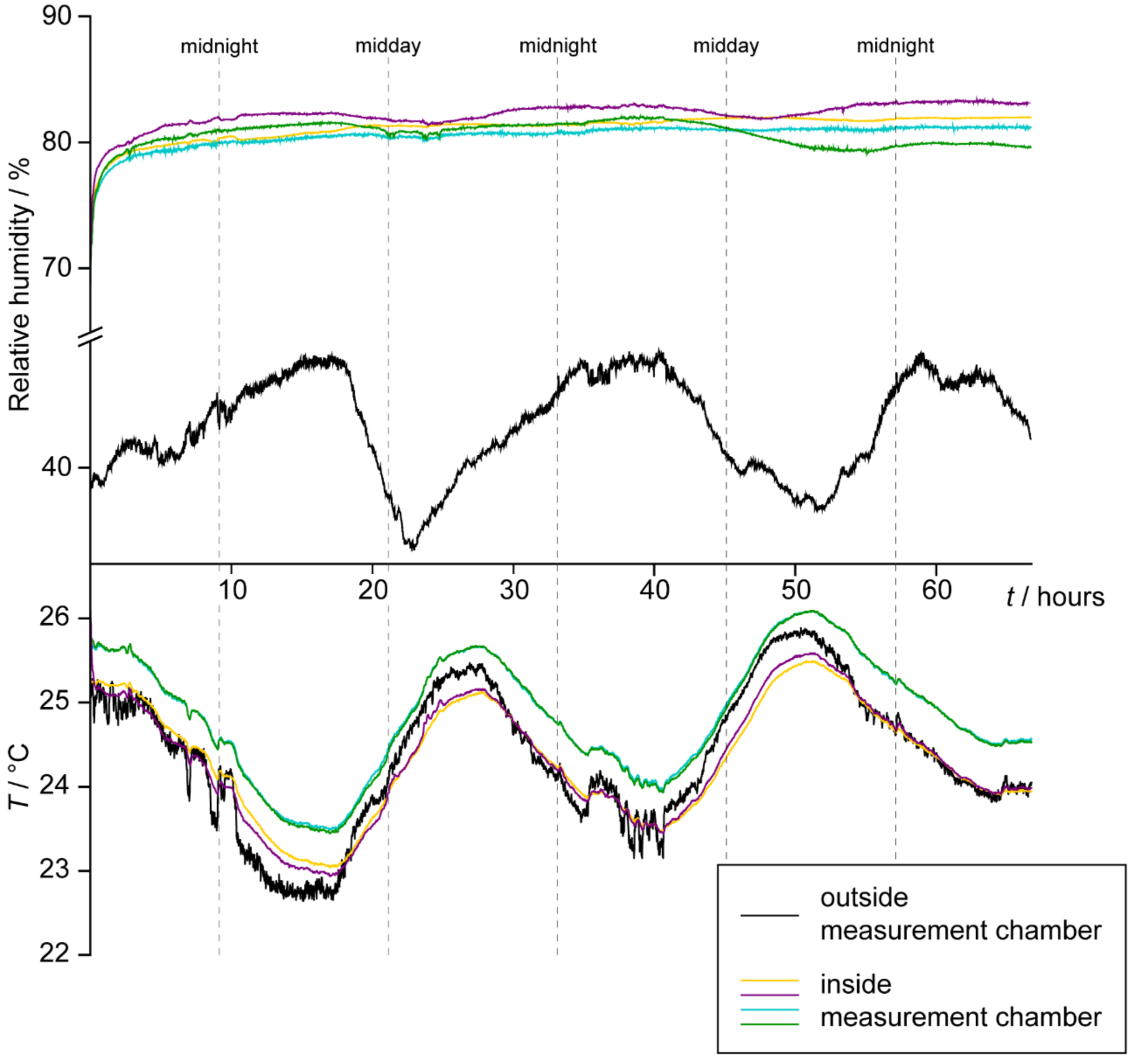


**Figure S1**. The relative humidity (top) and temperature (bottom) measured inside (yellow, purple, teal and green lines) and outside (black line) the measurement chamber, demonstrating the stability of the internal relative humidity despite changing temperature and external relative humidity. Each internal measurement was performed in an individual measurement chamber. A standard salt uptake experiment with kale seedlings was performed during these measurements, where 30 μL 0.1 M KCl was added through the measurement port at least 2 hours after measurements began; no obvious change in humidity was detected during this treatment step.

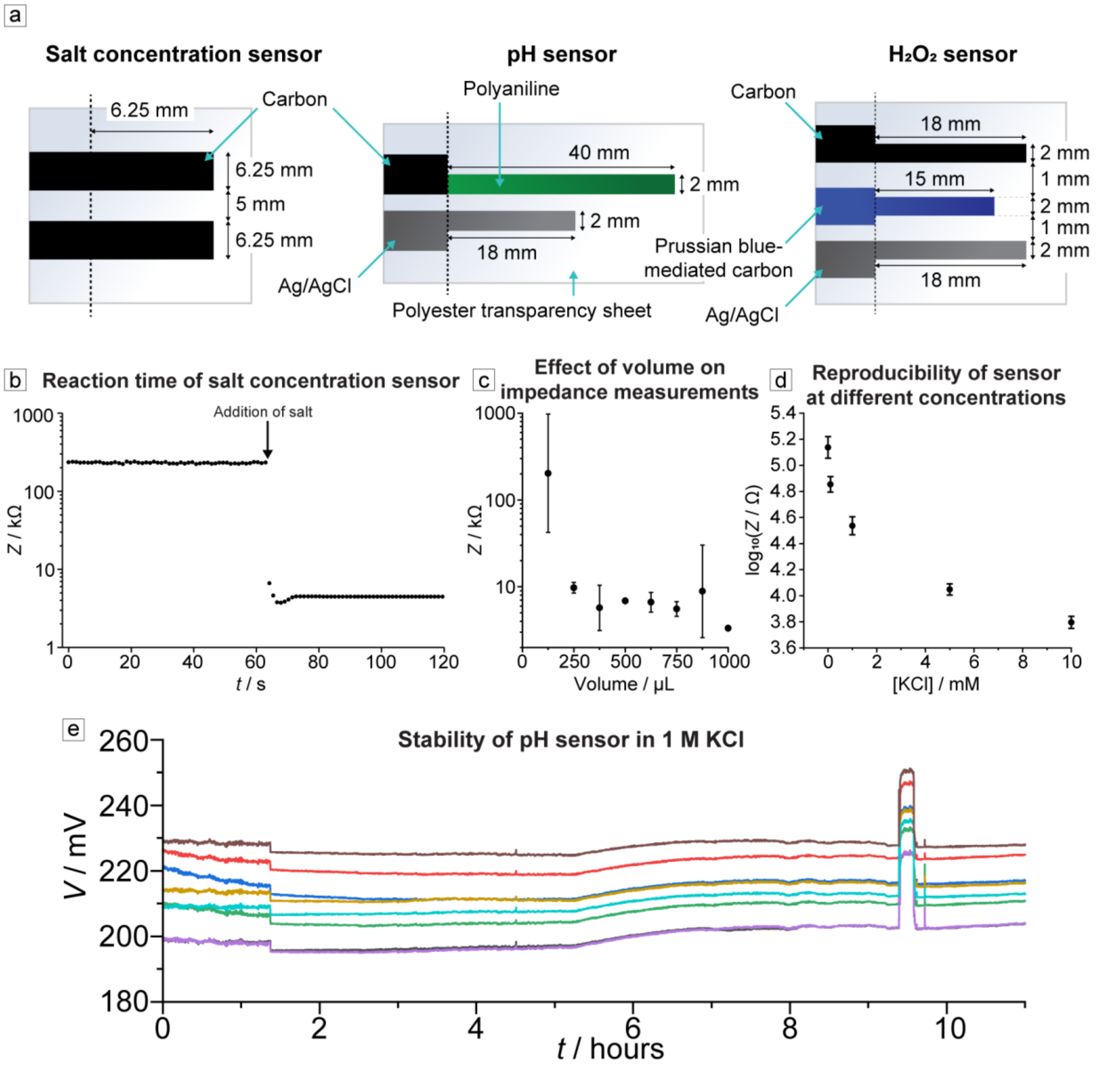


**Figure S2. a)** Dimensions and materials of our three sensors; **b)** Sensor response to sudden change in salt concentration. Arrow indicates time of addition of KCl (0.1 M, 30 μL) to paper disc containing 270 μL deionized water; **c)** Effect of volume of KCl (0.1 M) on impedance measurements. Error bars indicate 1 standard deviation (n = 2); **d)** Reproducibility of salt sensors at different concentrations. The logarithm of impedance of printed sensor with 500 μL solution in paper disc, at KCl concentrations of 0, 0.1, 1, 5 and 10 mM. Error bars show standard deviation (n = 5); e**)** Stability of our pH sensor in 1 M KCl.

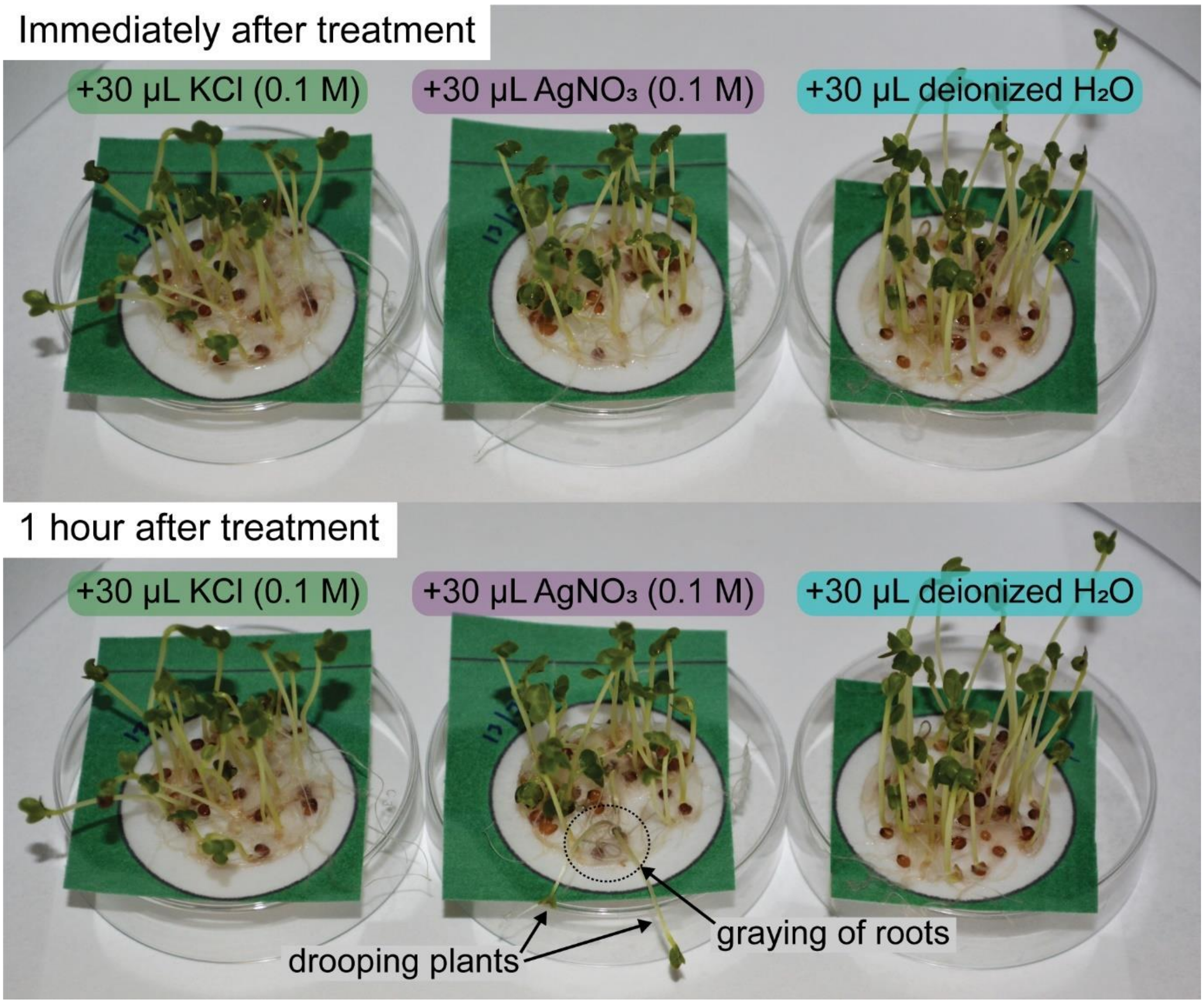


**Figure S3.** Treating kale seedlings with $AgNO_3$ solution resulted in metallic silver precipitating onto the roots, leaving a gray coloration. Some drooping of the plants was also observed for those plants where the gray coloration was the most obvious. No color change was observed with treatment of KCl or water.

**Table S1.** The constants A and n and adjusted R-squared value of the line of best fit from plotting log(*Z*) against log(*c*), where the relationship of electrical impedance (*Z*) and salt concentration (*c*) had the form $Z=Ac^n$.

| **Salt** | **A** | **n** | **Adj. R-squared** | **Linear range found** | **Number of measurements in linear range** |
|---|---|---|---|---|---|
| $AgNO_3$ | 304.1865 | -0.65131 | 0.80214 | 0.1 mM–0.1 M | 15 |
| $BaCl_2$ | 142.8828 | -0.64357 | 0.80206 | 0.125–1 mM | 8 |
| $Ca(NO_3)_2$ | 258.6128 | -0.59552 | 0.767 | 0.125–1 mM | 8 |
| $CaCl_2$ | 2.46082 | -1.32354 | 0.87006 | 0.125–1 mM | 8 |
| $Cd(NO_3)_2$ | 163.7948 | -0.63987 | 0.90484 | 0.125–1 mM | 8 |
| $CdCl_2$ | 334.349 | -0.56173 | 0.85033 | 0.125–1 mM | 8 |
| $CdSO_4$ | 6704.401 | -0.37509 | 0.90133 | 0.125–1 mM | 8 |
| $Cu(NO_3)_2$ | 118.3123 | -0.67205 | 0.98613 | 0.125–1 mM | 8 |
| $CuCl_2$ | 286.7742 | -0.5752 | 0.971 | 0.1 mM–0.1 M | 15 |
| $CuSO_4$ | 383.0893 | -0.53075 | 0.98461 | 0.125–1 mM | 8 |
| $GdCl_3$ | 433.6107 | -0.48196 | 0.76721 | 0.125–1 mM | 8 |
| $K_2SO_4$ | 5.92611 | -1.0773 | 0.93764 | 0.125–1 mM | 8 |
| KCl | 291.1052 | -0.62373 | 0.93281 | 0.1 mM–0.1 M | 15 |
| $KH_2PO_4$ | 363.5465 | -0.62948 | 0.93313 | 0.1 mM–0.1 M | 12 |
| $KNO_3$ | 329.9286 | -0.58908 | 0.80775 | 0.1 mM–0.1 M | 15 |
| $LaCl_3$ | 148.6004 | -0.6505 | 0.8656 | 0.125–1 mM | 8 |
| $Mg(NO_3)_2$ | 580.50471 | -0.4769 | 0.85103 | 0.125–1 mM | 8 |
| $MgCl_2$ | 85.29233 | -0.70649 | 0.96773 | 0.125–1 mM | 8 |
| $Na_2CO_3$ | 429.32078 | -0.51237 | 0.82423 | 0.125–1 mM | 8 |
| $Na_2SO_4$ | 204.9085 | -0.63517 | 0.92592 | 0.125–1 mM | 8 |

| | | | | | |
|---|---|---|---|---|---|
| NaCl | 214.9661 | -0.69028 | 0.90808 | 0.1 mM–0.1 M | 23 |
| $NaH_2PO_4$ | 420.5329 | -0.60364 | 0.93292 | 0.1 mM–0.1 M | 12 |
| $NaNO_3$ | 492.7765 | -0.59425 | 0.92538 | 0.1 mM–0.1 M | 12 |
| NaOH | 0.11724 | -1.59736 | 0.84581 | 0.125–1 mM | 8 |
| $NH_4Cl$ | 339.8365 | -0.57598 | 0.886 | 0.1 mM–0.1 M | 12 |
| $NH_4H_2PO_4$ | 448.3632 | -0.61392 | 0.96546 | 0.1 mM–0.1 M | 12 |
| $NH_4NO_3$ | 185.2593 | -0.64442 | 0.78426 | 0.1 mM–0.1 M | 12 |
| $Ni(NO_3)_2$ | 187.17723 | -0.60573 | 0.86894 | 0.125–1 mM | 8 |
| $NiCl_2$ | 187.5858 | -0.61078 | 0.83476 | 0.125–1 mM | 8 |
